# Contrasting functions of ATP hydrolysis by MDA5 and LGP2 in viral RNA sensing

**DOI:** 10.1101/2023.05.25.542247

**Authors:** Rahul Singh, Yuan Wu, Alba Herrero del Valle, Kendra E. Leigh, Mark T. K. Cheng, Brian J. Ferguson, Yorgo Modis

**Affiliations:** Molecular Immunity Unit, Department of Medicine, University of Cambridge, MRC Laboratory of Molecular Biology, Cambridge CB2 0QH, UK; Cambridge Institute of Therapeutic Immunology & Infectious Disease (CITIID), Department of Medicine, University of Cambridge, Cambridge CB2 0AW, UK; Department of Pathology, University of Cambridge, Cambridge CB2 1QP, UK

## Abstract

Cytosolic long double-stranded RNA (dsRNA), among the most potent proinflammatory signals, is recognized by MDA5. MDA5 binds dsRNA cooperatively, forming helical filaments. ATP hydrolysis by MDA5 fulfills a proofreading function by promoting dissociation of shorter endogenous dsRNAs from MDA5 while allowing longer viral dsRNAs to remain bound leading to activation of interferon-β responses. Here, we show that adjacent MDA5 subunits in MDA5-dsRNA filaments hydrolyze ATP cooperatively, inducing cooperative filament disassembly. This amplifies the RNA footprint expansion that accompanies each round of ATP hydrolysis and allows MDA5 to displace tightly bound proteins from dsRNA. Our electron microscopy and biochemical assays show that LGP2 binds to dsRNA at internal binding sites through noncooperative ATP hydrolysis. Unlike MDA5, LGP2 has low nucleic acid selectivity and can hydrolyze GTP and CTP as well as ATP. Binding of LGP2 to dsRNA promotes nucleation of MDA5 filament assembly resulting in shorter filaments. Molecular modeling of the MDA5-LGP2 interface suggests that MDA5 interacts with dsRNA stem-bound rather than end-bound LGP2. We conclude that NTPase-dependent binding of LGP2 to internal sites on dsRNA increases the number and signaling output of MDA5-dsRNA complexes. Our work identifies novel molecular mechanisms contributing the selectivity and sensitivity of cytosolic dsRNA sensing.

**KEY POINTS:**

- Cooperative ATP hydrolysis in MDA5 filaments confers selectivity for dsRNA and displaces other proteins from RNA
- Noncooperative NTP hydrolysis by LGP2 induces binding to internal RNA sites with low selectivity
- RNA stem-bound LGP2 nucleates assembly of MDA5 signaling complexes on a broader set of RNA ligands

## INTRODUCTION

Viruses deliver or generate RNA in the host cell cytosol for viral gene expression by cellular machinery. When present in the cytosol, double-stranded RNAs (dsRNAs) are among the most potent proinflammatory pathogen associated molecular patterns (PAMPs) (1). The antiviral innate immune responses induced by dsRNA sensors must be sensitive enough to detect infection and specific enough to avoid activation by cellular RNA. Failure to balance sensitivity with specificity can lead to viral proliferation or autoinflammatory disease (2). The primary sensors of cytosolic dsRNA in vertebrates are RIG-I (3,4), MDA5 (4–6), LGP2 (7–11), often referred to as the RIG-I-like receptors (RLRs), and protein kinase R (12,13). These are complemented in mammals by the oligoadenylate synthase (OAS) family of proteins (14,15) as well as ZBP1, and in humans, NLRP1 (16). dsRNAs longer than one hundred base pairs (bp) are recognized by MDA5 (5,6). MDA5 binds dsRNA cooperatively, forming helical filaments with its RIG-I-like superfamily 2 helicase module and C-terminal domain (CTD) (17–22). MDA5 filament formation induces the tandem caspase recruitment domains (CARDs) of MDA5 to form oligomers (17,18). These MDA5 CARD oligomers recruit MAVS via CARD-CARD interactions, nucleating the assembly of MAVS CARD into microfibrils (23,24), which activate potent interferon-β and NF-κB inflammatory responses (5,6,23).

MDA5 is the primary innate immune sensor for many viruses (5,6), including SARS-CoV-2 (25,26), but efficient proofreading and finely tuned signal transduction are necessary for a measured response to infection. Structural studies have shown that ATP hydrolysis by MDA5 performs a mechanical proofreading function (21,22). The ATPase cycle is coupled to conformational changes in MDA5-dsRNA filaments that test the interactions of MDA5 with the bound RNA, promoting dissociation of MDA5 from endogenous RΝΑs with shorter or imperfect double-stranded regions, while allowing MDA5 to remain bound to longer virus-like dsRNAs long enough to activate signaling (21,22). A-to-I deamination of dsRNA by ADAR1, which weakens base pairing and causes bulges, is nevertheless required to prevent MDA5-dependent autoinflammatory signaling induced by endogenous RNAs with high double-stranded content, for example annealed inverted Alu repeats (27–30). MDA5 mutations that disrupt proofreading, for example by inhibiting ATP hydrolysis, or otherwise increase baseline interferon signaling have been linked to autoinflammatory disorders in heterozygous human patients, including Aicardi-Goutières syndrome (AGS), Singleton-Merten syndrome (SMS), spastic-dystonic syndrome, and neuroregression (31–33).

The conformational changes in MDA5-dsRNA filaments associated with ATP hydrolysis appear to fulfill other functions in addition to proofreading. ATP hydrolysis causes a 1-bp expansion of the RNA binding footprint of MDA5 from 14 to 15 base pairs (21). If adjacent MDA5 subunits in a filament hydrolyze ATP in a sequential or cooperative manner, this RNA footprint expansion would be amplified. Expansion of the dsRNA footprint could explain the reported capacity of MDA5 to repair filament discontinuities and displace viral proteins from dsRNA in an ATPase-dependent, CARD-independent manner (34,35). Indeed, ATP hydrolysis by RIG-I promotes assembly of signaling-active oligomers through local translocation, or threading of successive RIG-I monomers onto dsRNA ends (36,37). However, whether ATP hydrolysis in adjacent MDA5 filament subunits is cooperative and drives expansion of the RNA footprint of MDA5 remains to be determined.

Multiple cofactors further increase the sensitivity and selectivity of MDA5-RNA recognition. LGP2, arguably the most important MDA5 cofactor, promotes MDA5 filament formation on dsRNA (10,38–41). LGP2 contains a DExD/H-box helicase and CTD similar to those of MDA5 but lacks CARDs. Structural studies show that the LGP2 CTD can bind dsRNA blunt ends (8–10). LGP2 RNA end binding has been proposed to nucleate MDA5 oligomerization on short (or otherwise suboptimal) dsRNAs (10,42). LGP2 has also been found incorporated within MDA5 filaments, accelerating MDA5 filament formation and resulting in more numerous, shorter filaments that retain signaling activity (40,41). The ATPase activity of LGP2 has been reported to promote dissociation of MDA5 from RΝΑ (41), but a separate study reported that ATP hydrolysis by LGP2 enhances LGP2 RNA binding and hence promotes MDA5 signaling (39). It remains unclear, therefore, whether LGP2 interacts directly with MDA5, or exactly how the RNA binding and ATPase activities of LGP2 contribute to sensitive and selective dsRNA recognition by MDA5.

Here, we use a combination of electron microscopy and biochemical assays to show that adjacent MDA5 subunits hydrolyze ATP cooperatively within MDA5-dsRNA filaments and furthermore that RNA footprint expansion from this cooperative ATP hydrolysis within MDA5-RNA filaments can displace tightly bound proteins from dsRNA. We show that dsRNA-bound LGP2 promotes nucleation of MDA5 filament assembly, resulting in a greater number of shorter MDA5 filaments. At higher concentrations, LGP2 competes with MDA5 for dsRNA binding, indicating that LGP2 can bind dsRNA internally, as well as at the ends. Unlike MDA5, LGP2 has low nucleic acid selectivity and hydrolyzes CTP and GTP as well as ATP. Nucleotide hydrolysis by LGP2 is noncooperative and coupled to dsRNA binding at internal binding sites. Molecular modeling of the MDA5-LGP2 interface with AlphaFold suggests that LGP2 recruits MDA5 to internal sites on dsRNA rather than the dsRNA ends. Our work explains the ATP-dependent, interferon-independent antiviral activity of MDA5 and suggests that LGP2 amplifies MDA5 signaling by promoting the formation of stable MDA5 filaments.

## MATERIALS AND METHODS

### RNA and DNA reagents

RNAs were transcribed *in vitro* using the MEGAscript T7 Transcription Kit (Invitrogen, cat. no. AM1333) or HiScribe® T7 High Yield RNA Synthesis Kit (New England BioLabs, cat. no. E2040S) following the manufacturers’ protocols. The 40-bp ssRNA and 100-bp, 300-bp and 1-kb dsRNA (and ssRNA) contained the first 40, 100, 300 or 1000 bases of the mouse *IFIH1* gene, respectively. The complementary RNA strands were transcribed from DNA templates with a preceding 5′ TAATACGACTCACTATAG 3′ sequence. The *in vitro* transcription reactions were performed at 37°C for 2, 4, 6 h or overnight. Transcripts were treated with TURBO DNase and purified with the PureLink RNA Mini Kit (ThermoFisher, cat. no. 12183018A) or the Monarch RNA Cleanup Kit (500 μg) (New England BioLabs, cat. no. T2050L). Samples were eluted in a nuclease-free duplex annealing buffer: 30 mM HEPES pH 7.5, 0.1 M KCl (IDT). Eluted transcripts were incubated at 95°C for 5 min and cooled to room temperature over 2 h to eliminate secondary structure and enable annealing of complementary strands of RNA where necessary. To obtain internally biotinylated dsRNA, 10x Biotin RNA Labelling Mix (Roche, 11685597910), consisting of 10 mM each of ATP, CTP and GTP, 6.5 mM UTP and 3.5 mM Biotin-16-UTP, was added to the *in vitro* transcription reaction mix. 100 bp and 1-kb dsDNA was obtained by PCR amplification of the first 100 or 1000 bases of the mouse *IFIH1* gene. DNA:RNA hybrids were produced using *in vitro* transcribed ssRNA to generate and hybridize the complementary cDNA with the RevertAid™ H Minus Reverse Transcriptase (RT) kit (ThermoFisher, cat. no. EP045). Some samples were incubated with RNase H (New England BioLabs, cat. no. M0297S), DNA:RNA hybrid-specific nuclease, for 30 min at 37°C (see Figure 3A). High molecular weight poly(I:C) RNA was purchased from InvivoGen (cat. no. tlrl-pic). 4-kb Circular dsRNA was a gift from David Dulin (VU University Amsterdam). The ssRNAs with complex secondary structure were gifts from Irene Díaz-López from the laboratory of Venki Ramakrishnan (MRC Laboratory of Molecular Biology). See **Supplementary Data** for RNA sequences.

### Expression and purification of MDA5

A gene encoding murine MDA5 (*Ifih1,* UniProt: Q8R5F7) was cloned into the pET28a vector with an N-terminal hexa-histidine tag followed by a tobacco etch virus (TEV) protease cleavage site as described (18). MDA5 residues 646–663, in the flexible L2 surface loop of the helicase 2 insert domain (Hel2i), were deleted for solubility, resulting in a 114-kDa polypeptide chain. This ΔL2 loop deletion does not affect the dsRNA binding, ATPase or interferon signaling activities of MDA5 (18,20,21). Expression plasmids for MDA5 mutants were generated by site-directed mutagenesis.

*Escherichia coli (E. coli)* Rosetta2(DE3)pLysS cells (Novagen, cat. no. 71403) were transformed with pET28a-MDA5ΔL2-His_6_ and grown in 2xTY medium to OD_600_ 0.4-0.6 at 37°C. After cooling to 17°C, protein expression was induced with 0.5 mM isopropyl-β-D-1-thiogalactopyranoside (IPTG) overnight at 17°C. Harvested cells were resuspended in 30 mM HEPES pH 7.7, 0.15 M NaCl, 5% glycerol, 1 mM Tris(2-carboxyethyl)phosphine (TCEP), cOmplete™, EDTA-free Protease Inhibitor Cocktail (Roche, cat. no. 11873580001) and Benzonase® Nuclease (Merck, cat. no. 70746), then lysed by ultrasonication on ice. The lysate was centrifuged at 37,500 g for 1 h. The supernatant was loaded onto a pre-equilibrated 5 ml HisTrap HP column (Cytiva, cat. no. 17-5248-02), washed with 30 mM HEPES pH 7.7, 0.5 M NaCl, 20 mM imidazole and 1 mM TCEP, and MDA5 was eluted with 30 mM HEPES 7.5, 0.15 M NaCl, 5% Glycerol, 0.3 M imidazole and 1 mM TCEP. MDA5 was further purified on a Resource Q anion exchange column (Cytiva) (buffer A: 20 mM HEPES 7.5, 0.1 M NaCl, 2 mM dithiothreitol (DTT); buffer B: 20 mM HEPES 7.5, 1 M NaCl, 2mM DTT), and a Superdex 200 Increase 10/300 GL size-exclusion column (Cytiva) in 20 mM HEPES pH 7.8, 0.15 M KCl, 1 mM DTT, 5% glycerol. Purified protein was flash-frozen and stored at −80°C.

### Expression and purification of LGP2

A gene encoding murine LGP2 (*Dhx58*, UniProt: Q99J87) was cloned into the pACEBAC1 baculovirus transfer vector with an N-terminal 2x-Strep tag followed by TEV protease cleavage site. The K30G and R32G mutations were introduced into the LGP2 construct by site-directed mutagenesis.

The pACEBAC1-2xStrepTEV-LGP2 plasmids were transformed into *E. coli* DH10Bac cells. Purified bacmid DNA was transfected into Sf9 insect cells (Invitrogen) to produce secreted recombinant baculovirus. For LGP2 expression, Sf9 cells were infected at a density of 3 × 10^6^ cells/ml with 0.5% (v/v) of third passage (P3) baculovirus stock and incubated at 27°C. The cells were harvested 60-72 h post-infection. The cells were lysed with a chilled Dounce homogeniser in 30 mM HEPES, pH 7, 0.3 M NaCl, 1 mM TCEP, 5% glycerol supplemented with EDTA-free Complete Protease Inhibitor (Roche) and Salt-Active Nuclease (Sigma-Aldrich, cat. no. SRE0015). The lysate was clarified by centrifugation. The lysate supernatant was incubated on a nutator at 4°C for 2-3 h with 1 ml of Strep-Tactin Sepharose resin (IBA-Lifesciences, cat. no. 2-1201-025) equilibrated in 30 mM HEPES, pH 7, 0.15 M NaCl, 1 mM TCEP, 5% glycerol, and EDTA-free Complete Protease Inhibitor (Roche). The protein was eluted with 2.5 mM d-desthiobiotin pH 8. The eluate was further purified on a Superdex 200 Increase 10/300 GL size-exclusion column in 25 mM HEPES pH 7.7, 0.15 KCl, 1 mM TCEP.

### Negative stain electron microscopy

Carbon film in 300-mesh grids (Agar Scientific) were glow discharged at 25 mA for 1 min. Samples were applied to the grids and washed with RNase-free water before staining with 2% (w/v) uranyl acetate. To visualize MDA5 filaments from the cooperativity assays, protein was incubated on ice for 30 min with 10 mM ATP-γ-S with a final concentration of 200 nM MDA5 and 10 nM 1-kb dsRNA. To visualize LGP2-MDA5 complex formation, 73 nM MDA5 and 73, 146, or 730 nM LGP2 were incubated with 4 mM ATP and 4.4 nM 1-kb dsRNA on ice for 30 min. Samples containing MDA5 and 73, 146 and 730 nM LGP2 were rapidly diluted 1:2, 1:3 and 1:10 fold, respectively, before staining. All negative stain EM images were collected on a 120 kV Tecnai G2 Spirit TEM (ThermoFisher). Images were collected at −2 to −4 µm defocus, 26,000×magnification, and 4 Å per pixel. Results were analyzed and filament length quantified with ImageJ (v1.5).

### ATPase assays

ATPase activities were measured with the ATPase/GTPase Activity Assay Kit (Sigma-Aldrich, cat. no. MAK113) based on the colorimetric change of malachite green in the presence of inorganic phosphate produced by ATP hydrolysis. Reactions in clear, flat-bottom 96-well plates were read using a Clariostar microplate reader (BMG Labtech, Germany) at 620 nm. Reactions contained 180 nM protein, 25 ng nucleic acid, 4 mM ATP and 4 mM MgSO_4_ unless stated otherwise. Reactions were incubated at 37°C for 3 mins and quenched with EDTA. For the cooperativity assays, wild-type and ATPase-deficient variants were mixed at various molar ratios while maintaining the same total protein concentration of 200 nM. The WT:mutant mixtures were incubated with 1.25 nM 1-kb dsRNA at 37°C for 3 min. Results were analyzed with Prism v9.5.1 (GraphPad).

### Biotin-Streptavidin Displacement Assay

Biotinylated RNA duplexes were generated by incorporation of Biotin-16-UTP (Sigma-Aldrich) during *in vitro* transcription of 1-kb sense and antisense RNA strands (see above for *in vitro* transcription protocol). Ratios of incorporation were calculated to synthesize RNA duplexes with an average of 2 to 8 conjugated biotins. 2 μg/μl biotinylated dsRNA and unlabeled control were incubated with 55 mM streptavidin (ThermoFisher, cat. no. 211122) dissolved in nuclease-free water with 20 mM potassium phosphate pH 6.5. This mixture was incubated with 350 nM mMDA5, 245 mM biotin (Sigma-Aldrich, B4501; from a stock solution in 0.5% DMSO and DEPC water) and 10 mM ATP-MgSO_4_ for 30 min at 37°C. Streptavidin displacement was visualized by gel electrophoresis on 1% agarose or 4-20% TBE polyacrylamide gels stained with SYBR™ Gold Nucleic Acid Gel Stain (ThermoFisher, cat. no. S11494). The reactions were incubated at 65°C for 5 min in 0.5 M potassium phosphate pH 7.4, 0.5 M NaCl, 6% SDS, and 1x SDS Gel Loading Dye (New England BioLabs, cat. no. B7024S) – this treatment displaced MDA5 and LGP2 but not streptavidin from biotinylated RNA. The samples were loaded on the gels and the gels were run in TBE buffer at 200 V and 4°C. Gels were stained with DNA stain at room temperature for 2 h. Gels were imaged in a G:BOX imaging system and Gene Sys v1.5 (Syngene).

### Bilayer interferometry (BLI) protein-RNA binding assays

Pierce™ RNA 3’ Ed Biotinylation Kit (ThermoFisher, cat. no. 20160) was used to label the 3’ end of 1-kb RNA duplexes (produced as described above) with biotin. The 3’-biotinylated RNA was immobilized on an Octet® streptavidin (SA) Biosensor (Sartorius). 3 μg of biotinylated RNA were incubated per sensor for 300 s. MDA5 and LGP2 were added, individually or together, at 0.1 µm or 1 μM. The optical interference pattern of white light was used to measure binding kinetics with the Octet® BLI Label-Free Detection System (Sartorius). MDA5 and LGP2 dissociation was measured by flowing buffer (20 mM HEPES pH 7.4, 0.15 M KCl and 1 mM TCEP) over the sensor. BLI kinetic data was analyzed with Octet® Analysis Studio (Sartorius) to determine dissociation constants (*K_d_)*. *K_d_* values were calculated from the association rate.

### Electrophoretic mobility shift assays (EMSA)

Mixtures containing 100 ng nucleic acid and 1 μM MDA5 or LGP2 were incubated for 15 min at 4°C in 20 mM HEPES pH 7.4, 0.15 M KCl and 1 mM TCEP. After addition of NativePAGE™ Sample Buffer (ThermoFisher, cat. no. BN2003), samples were loaded onto a 4-20% polyacrylamide gel and run in in Tris-borate (TB) buffer (0.1 M Tris, 0.1 M Boric acid, pH 8.3) for 90 min at 200 V at 4°C. Gels were stained with SYBR™ Gold Nucleic Acid Gel Stain (ThermoFisher, cat. no. S11494) and imaged in a G:BOX imaging system and Gene Sys v1.5 (Syngene).

### AlphaFold2 modeling

Atomic models of MDA5-LGP2 complexes were predicted with a local installation of AlphaFold2-v2.3.1 (43,44) using multimer mode as implemented by ColabFold v1.5.2 (45) with the use of MMseqs2 for multiple sequence alignment (MSA). The amino acid sequences of human or mouse LGP2 (all residues) and MDA5-ΔCARD (residues 306-1025) were used as the input. Polypeptide geometry in the models was regularized by post-prediction relaxation using the Amber force field. The maximum number of recycles was set to 20. No MSA cutoff was used. The sequence pairing mode was set to pair sequences from the same species and unpaired MSA. The multimer model was set to multimer-v3. Use of available template files from the Protein Data Bank had no significant effect on atomic model prediction. Five atomic models were output from each AlphaFold run. Protein interfaces were analyzed with the Protein interfaces, surfaces, and assemblies (PISA) service at the European Bioinformatics Institute [http://www.ebi.ac.uk/pdbe/prot_int/pistart.html] (46). Atomic coordinates of a model generated by AlphaFold-Multimer containing two human MDA5 molecules and one human LGP2 molecule are available as a separate **Supplementary Data** file.

### Statistics

No statistical methods were used to predetermine sample size, experiments were not randomized, and the investigators were not blinded to experimental outcomes. ATPase assays were performed at least three times in independent experiments. Scatter plots, histograms and error bars were plotted with GraphPad Prism 9.5.1 and Microsoft Excel v.16.72.

## RESULTS

### Adjacent subunits in MDA5-dsRNA filaments hydrolyze ATP cooperatively

The ATPase activity of MDA5 has been reported to fulfill additional functions other than proofreading, including repair of filament discontinuities and displacement of viral proteins from dsRNA (34,35), but the molecular basis for these functions remains unknown. ATP hydrolysis causes a 1-bp expansion of the RNA binding footprint of MDA5 (21) so we hypothesized that adjacent MDA5 subunits in a filament may hydrolyze ATP sequentially or cooperatively and hence amplify this RNA footprint expansion sufficiently to close gaps in filaments and displace other RNA-bound proteins. To test this hypothesis, we measured the ATPase activity of mouse MDA5-dsRNA filaments containing increasing proportions of the ATPase deficient MDA5 R338G mutant (corresponding to R337G in humans (32)), which has 8% of wild-type ATPase activity but can still bind and form filaments on dsRNA (**Figure 1A-B, Supplementary Figure S1**). An analogous approach was used to detect sequential ATP hydrolysis in the hexameric ATPase FtsK (47). If MDA5 subunits hydrolyzed ATP independently, the loss of filament ATPase activity would be expected to be proportional to the amount of ATPase-deficient mutant incorporated. Instead, the ATPase activity dropped exponentially with increasing amounts of ATPase-deficient mutant, with MDA5-dsRNA filaments containing 20% of the ATPase-deficient mutant having less than half the ATPase activity of wild-type MDA5 filaments (**Figure 1A**). The ATPase activity of wild-type MDA5-dsRNA complexes also increased exponentially as a function of increasing MDA5 concentration, reflecting the cooperativity of both filament assembly and ATP hydrolysis (**Figure 1B**). In contrast, the ATPase activity of an MDA5 mutant with mutations that prevent filament formation, L397A/K398A/I399A (21), was directly proportional to the concentration of the MDA5 mutant, indicating noncooperative ATP hydrolysis (**Figure 1B**). The ATPase activity of apyrase, which is monomeric in solution, was measured to provide a reference curve for noncooperative ATPase activity (**Figure 1C**). Together, these data indicate that ATP hydrolysis in wild-type MDA5 filaments is cooperative, and that this cooperativity depends on filament formation.

**Figure 1.**
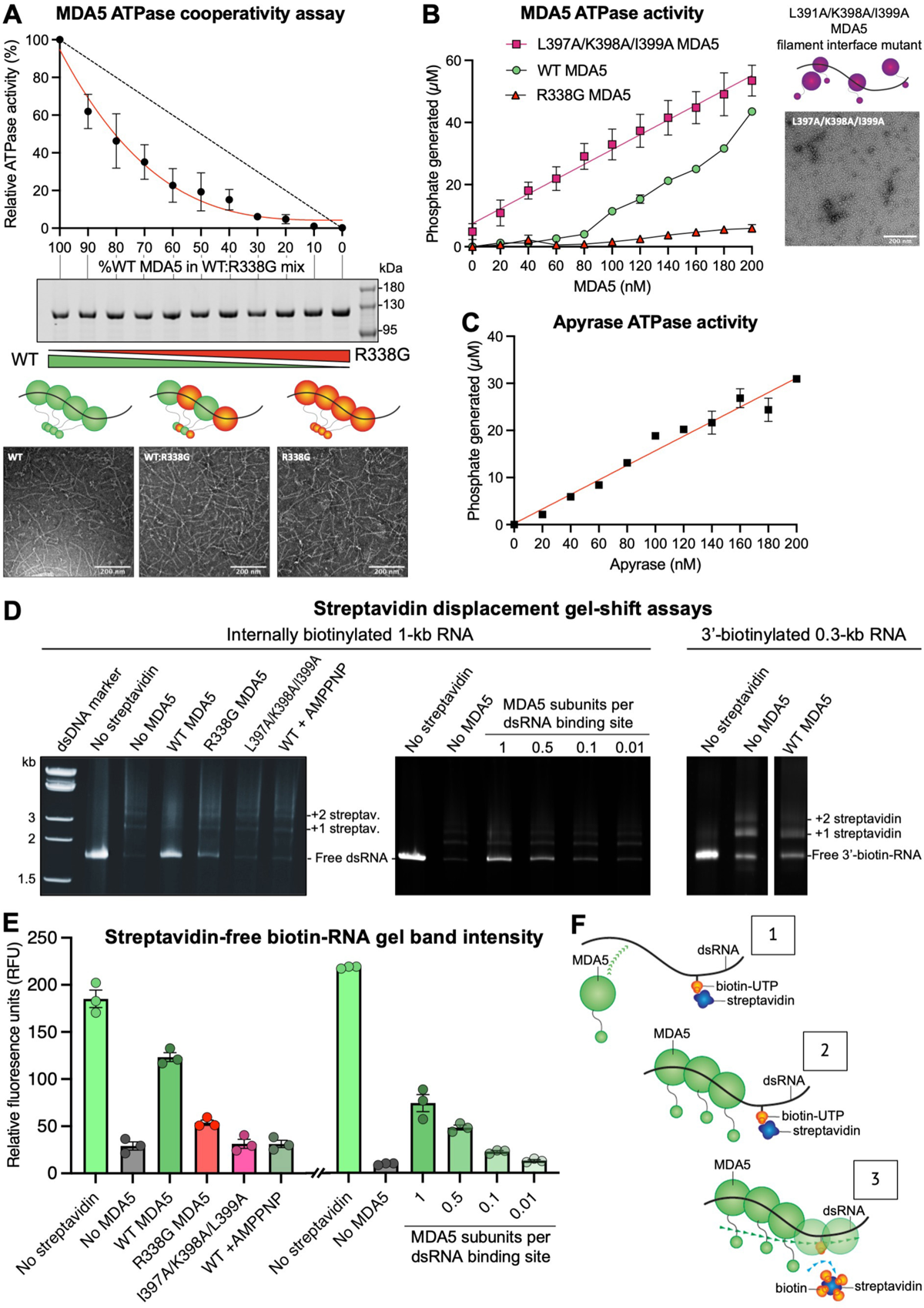
ATPase cooperativity and streptavidin displacement assays for MDA5-dsRNA filaments. (A) ATPase activity of mouse MDA5-dsRNA filaments containing increasing proportions of the R338G MDA5 variant, which is ATPase deficient (see **Supplementary Figure S1A**). Dashed line, the linear relationship expected in the absence of cooperativity. Middle panel: Coomassie-stained SDS-PAGE gel showing the total protein in each reaction mixture. Bottom panel: Negative-stain electron micrographs of WT MDA5, a 1:1 WT:R338G MDA5 mix, and R338G MDA5 in the presence of 1-kb dsRNA. Scale bars, 200 nm. (B) Left, ATPase activity of MDA5-dsRNA complexes as a function of MDA5 concentration. Right, negative-stain electron micrograph of the filament interface mutant in the presence of 1-kb dsRNA. Scale bar, 200 nm. (C) ATPase activity of apyrase, a monomeric ATPase, as a function of apyrase concentration. (D) Polyacrylamide gel-shift assay for displacement of streptavidin from biotinylated dsRNA. Samples were incubated in semi-denaturing conditions prior to electrophoresis (See Materials and Methods) to displace MDA5 but not streptavidin from the RNA. One MDA5 binding site was defined as 15 bp of dsRNA. (E) Quantification of streptavidin-free biotinylated RNA from gel band densitometry. Error bars show the s.e.m. of three independent experiments (n = 3) including the gels shown in (D). See **Supplementary Data** for source data. (F) Proposed mechanism of streptavidin displacement from biotinylated RNA by cooperative ATP hydrolysis within MDA5 filaments.

### MDA5 ATP hydrolysis displaces streptavidin from biotinylated dsRNA

A predicted consequence of cooperative ATP hydrolysis in MDA5-dsRNA filaments is that adjacent MDA5 subunits will hydrolyze ATP sequentially, causing a wave of ATP hydrolysis to spread along the filament or a segment thereof. This process could amplify the 1-bp RNA footprint expansion that accompanies each round of ATP hydrolysis and hence potentially displace any proteins bound to the RNA. To test this hypothesis, we generated 1-kb dsRNA containing a small fraction of internally incorporated or 3’-end incorporated biotinylated nucleotides, allowed streptavidin tetramers to bind to the biotin moieties, and asked whether MDA5 could displace the streptavidin from the RNA. Similar approaches were used previously to detect translocation of RIG-I and other helicases along nucleic acid duplexes (36,48). We found that MDA5 could displace streptavidin from internally biotinylated dsRNA in the presence of ATP (**Figure 1D-E**).

No streptavidin displacement was observed from 3’-end biotinylated dsRNA, or when the non-hydrolyzable ATP analog AMPPNP was used in place of ATP (**Figure 1D**). Efficient streptavidin displacement required a molar excess of MDA5 relative to the number of 15-bp dsRNA binding sites (**Figure 1D-E**), consistent with limited filament formation and ATP hydrolysis at small MDA5 to RNA molar ratios (**Figure 1B**). The monomeric, ATPase-active MDA5 mutant L397A/K398A/I399A was unable to displace streptavidin, confirming that streptavidin displacement requires both filament formation and ATP hydrolysis. The ATPase-deficient MDA5 mutant R338G had low but detectable streptavidin displacement activity, consistent with the residual ATPase activity of this mutant (8% of wild-type MDA5). We conclude that cooperative ATP hydrolysis allows MDA5 filaments to displace tightly bound proteins from dsRNA by amplifying the RNA footprint expansion that accompanies each round of ATP hydrolysis by a single filament subunit (**Figure 1F**).

### ATP hydrolysis by LGP2 is noncooperative and required for internal dsRNA binding

ATP hydrolysis by LGP2 had been reported to enhance binding of LGP2 to RNA and thus promote MDA5 signaling (39), but a subsequent study reported that the ATPase activity of LGP2 promoted dissociation of MDA5 from RΝΑ (41). We applied negative-stain electron microscopy (ns-EM) and biochemical assays to clarify the role of ATP hydrolysis by LGP2 in RNA binding and MDA5 filament formation. Ns-EM of LGP2 and dsRNA in the absence of MDA5 clearly showed that LGP2 can bind to internal dsRNA binding sites (**Figure 2A**). Circular dsRNA induced LGP2 to hydrolyze ATP at the same rate as linear dsRNA, demonstrating that the absence of RNA ends does not impede the nucleic acid-dependent ATPase activity of LGP2 (**Figure 3A**). Moreover, the LGP2 mutant R32G, which has 4% of wild-type LGP2 ATPase activity, was partially deficient for internal RNA binding (**Figure 2A**, **Supplementary Figure S1A**). LGP2 mutant K30G, which lacks ATPase activity, failed to bind RNA internally. Consistent with this, bilayer interferometry data showed that the LGP2 K30G mutant had a 20-fold lower binding affinity for 1-kb dsRNA than wild-type LGP2, with a *K_d_* of 1.3 µM for K30G versus 70 nM for wild-type (**Figure 2B**). Similar differences in relative binding affinities were observed with a 300-bp dsRNA and a 40-nt ssRNA, with lower binding affinities for the LGP2 mutants as expected (**Supplementary Figure S1B**). We detected no basal ATPase activity in LGP2 in the absence of nucleic acids (**Figure 3A**), in contrast to a previous study (39), but consistent with two other studies (10,41). Together our data show that ATP hydrolysis by LGP2 promotes dsRNA binding and specifically is required for stem binding to internal dsRNA sites.

**Figure 2.**
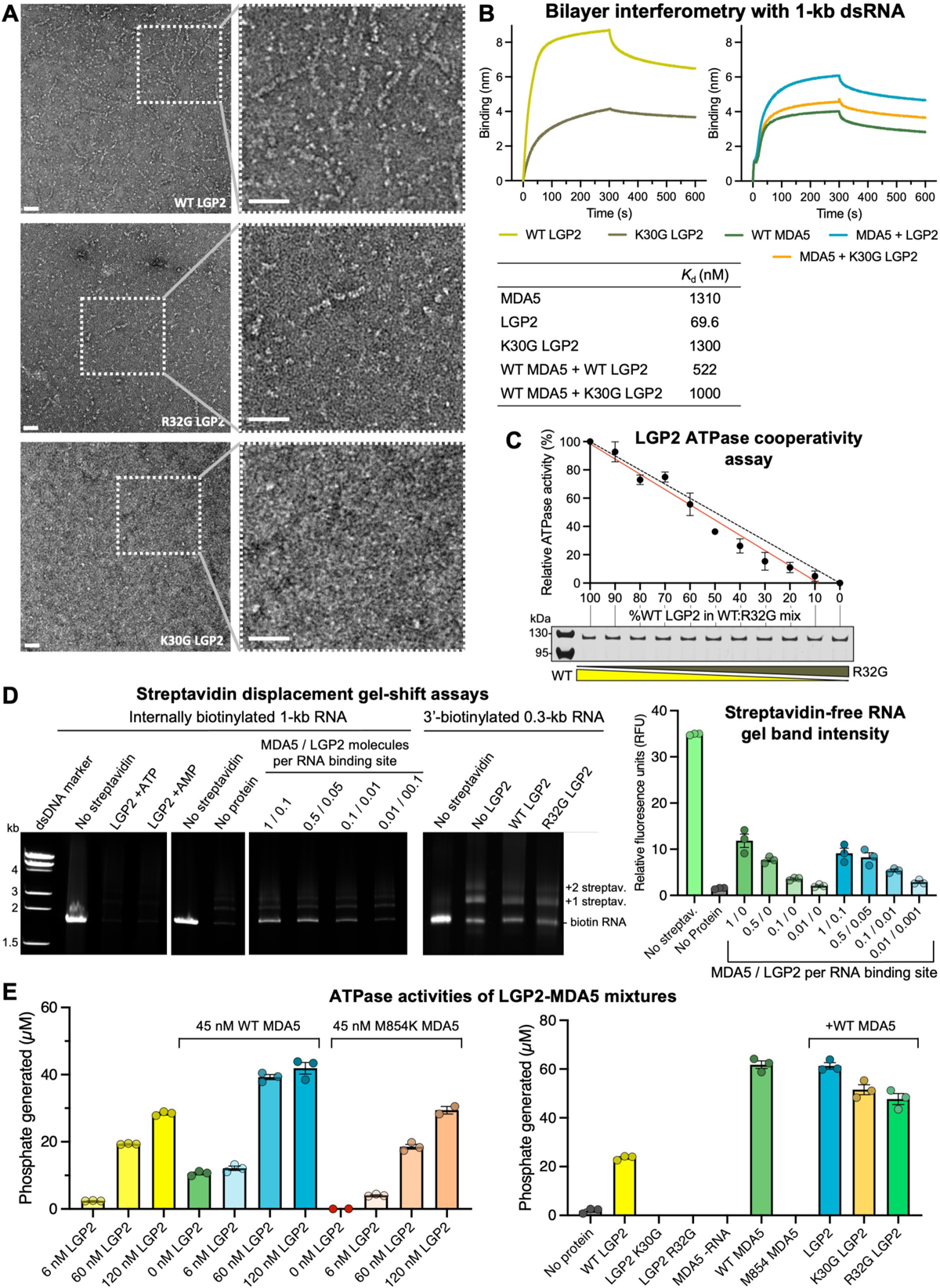
RNA binding, ATPase cooperativity and streptavidin displacement assays with LGP2 and LGP2-MDA5 mixtures. (A) Negative-stain electron micrographs of LGP2 (WT, R32G, and K30G variants) with 1-kb dsRNA. Scale bars, 50 nm. (B) Bilayer interferometry with 1-kb 3’-biotinylated dsRNA immobilized on a streptavidin biosensor and MDA5, LGP2 or MDA5-LGP2 mixtures in the mobile phase. Binding curves with immobilized 300-bp dsRNA and 40-nt ssRNA are shown in **Supplementary Figure S1B**. Dissociation constants derived from the curves are tabulated below. (C) ATPase activity of mouse LGP2-dsRNA filaments containing increasing proportions of the R32G LGP2 variant, which is ATPase deficient (see **Supplementary Figure S1A**). Lower panel: Coomassie-stained SDS-PAGE gel showing the total protein in each reaction mixture. (D) Polyacrylamide gel-shift assay for displacement of streptavidin from biotinylated dsRNA. Samples were incubated in semi-denaturing conditions (See Materials and Methods) to displace MDA5 and LGP2 but not streptavidin from the RNA. One binding site was defined as 15 bp of dsRNA. Right panel, quantification of streptavidin-free biotinylated RNA from gel band densitometry. The first six bars are also shown in Figure 1E. Error bars show the s.e.m. of three independent experiments (n = 3) including the gels shown on the left. See **Supplementary Data** for source data. (E) ATPase activities assay of different variants of LGP2 and MDA5 and their mixtures, with dsRNA binding sites in excess. The concentration of 15-bp dsRNA binding sites was 313 nM. Error bars show the s.e.m. of three independent experiments.

The ATPase activity of LGP2-dsRNA complexes containing increasing proportions of the ATPase-deficient R32G mutant decreased proportionally to the amount of the mutant present, indicating that ATP hydrolysis by LGP2 is noncooperative, in contrast to MDA5 (**Figure 2C**). Accordingly, LGP2 failed to displace streptavidin from internally biotinylated dsRNA or 3’-end biotinylated dsRNA (**Figure 2D**). Moreover, streptavidin displacement by MDA5 was unaffected by the addition of LGP2, implying that cooperative ATP hydrolysis by MDA5 filaments was unaffected by LGP2 (**Figure 2D**). The net ATPase activities of different mixtures of wild-type LGP2, LGP2 K30G, or LGP2 R32G and MDA5, with dsRNA in excess, were furthermore consistent with the ATPase activities of MDA5 and LGP2 being independent and additive, rather than cooperative (**Figure 2E**).

### LGP2 has low selectivity for nucleic acids and is a semi-selective NTPase

LGP2 is required to induce MDA5 signaling from RNAs with short or imperfect duplex regions, such as endogenous RNAs that escape editing by ADAR1 (42), suggesting that LGP2 has a broader dsRNA binding specificity than MDA5. Consistent with this, we found that 100-bp dsRNA, and a 1-kb DNA:RNA hybrid induced similar levels of ATP hydrolysis by LGP2 as long (>1 kb) linear or circular dsRNA (**Figure 3A**). Moreover, 100-bp dsDNA, 1-kb dsDNA, and 1-kb ssRNA induced 75-80% of the LGP2 ATPase activity observed with dsRNA. In contrast, 1-kb dsRNA, 4-kb circular dsRNA, poly(I:C), and ssRNAs predicted to have complex secondary structure each induced similar levels of ATP hydrolysis by MDA5, although both MDA5 and LGP2 formed stable complexes with circular (or linear) dsRNA and DNA:RNA hybrids in gel shift assays (**Figure 3B**, **Supplementary Figure S2**). The apparent binding affinity of LGP2 for 1-kb dsRNA from bilayer interferometry was 20-fold higher than that of MDA5, with K_d_ values of 70 nM and 1.3 µM, respectively (**Figure 2B**).

**Figure 3.**
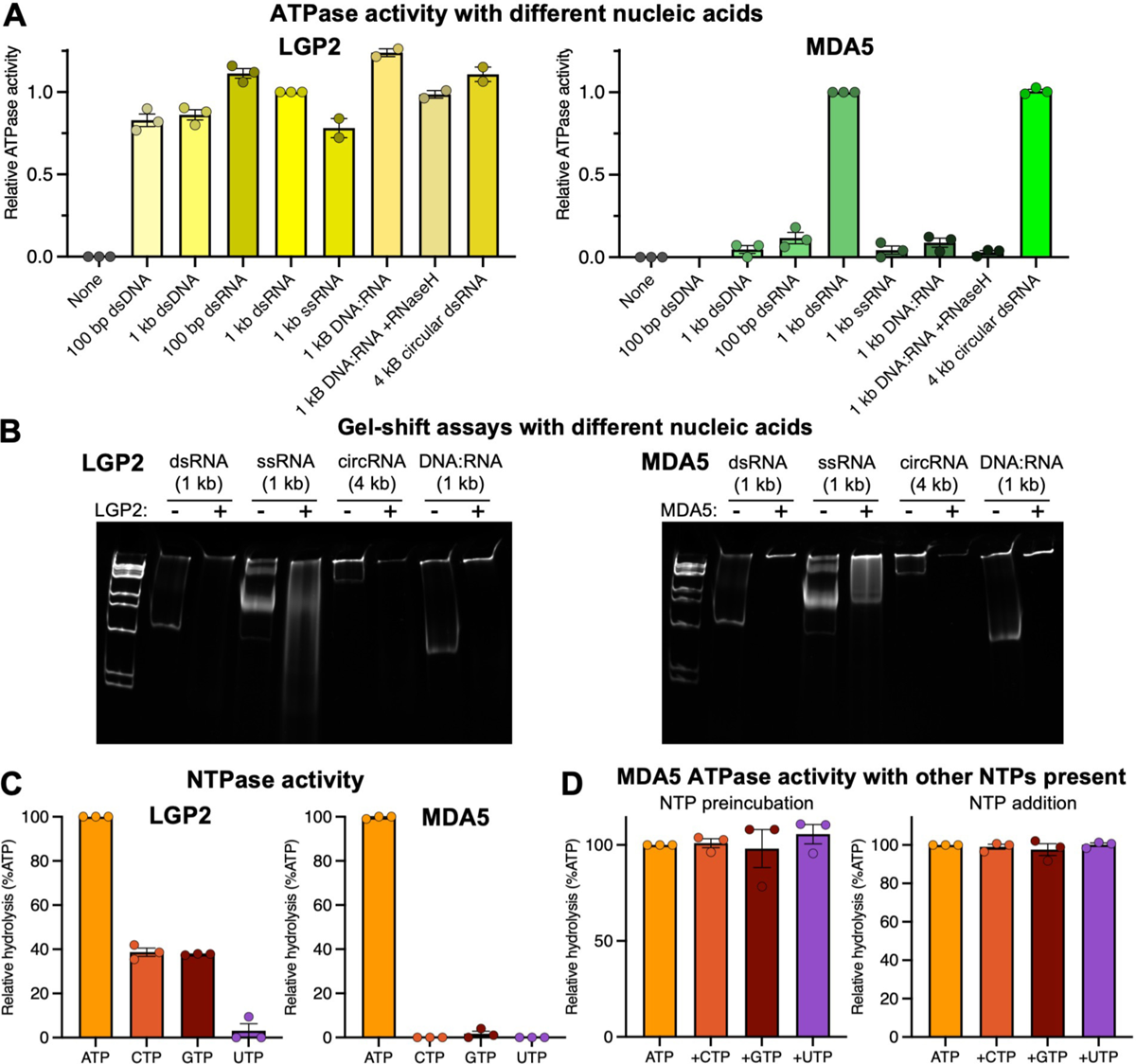
Nucleic acid and NTP selectivity for LGP2 and MDA5. (A) ATPase activity of LGP2 and MDA5 in the presence of different nucleic acid ligands, expressed as the relative activity normalized to the condition with 1-kb dsRNA. See **Supplementary Figure S2** for ATPase activity of MDA5 in the presence of additional ligands. Error bars show the s.e.m. of two or three independent experiments. (B) Gel-shift assay for binding of LGP2 and MDA5 to different nucleic acid ligands. 4-20% polyacrylamide gels were used with 1 kb DNA Ladder (New England BioLabs cat. no. N3232S). See **Supplementary Figure S2** for gel-shift assays with additional nucleic acid ligands. (C) Nucleotide hydrolysis activity of LGP2 and MDA5, expressed as percent of ATP hydrolysis activity. Error bars show the s.e.m. of three independent experiments. See **Supplementary Data** for source data. (D) ATPase activity of MDA5 following preincubation with or addition of CTP, GTP or UTP in the presence of ATP. ATPase activity is not affected by the other nucleotides, demonstrating that they do not compete with ATP. Error bars show the s.e.m. of three independent experiments.

Nucleotide hydrolysis assays showed that LGP2 is also less selective than MDA5 in its nucleotide utilization. LGP2 hydrolyzes CTP and GTP 40% as efficiently as ATP, in contrast to MDA5, which only hydrolyzes ATP (**Figure 3C**). No UTP hydrolysis could be detected for either LGP2 or MDA5. Addition of CTP, GTP or UTP did not affect MDA5 ATPase activity, demonstrating that the ATP binding pocket of MDA5 is selective for ATP (**Figure 3D**). Although DEAH/RHA and NS3/NPH-II helicases from superfamily 2 hydrolyze all four types of NTPs (49,50), the semi-selective NTP hydrolysis observed here for LGP2 is unexpected given that MDA5 is selective for ATP. We note that RNA viruses have unusual genomic nucleotide compositions (51) causing different NTPs to be consumed at different rates. This raises the question of whether LGP2 evolved its lower nucleotide selectivity as an adaptation to changes in the cellular nucleotide pool associated with infection.

### LGP2 promotes nucleation of MDA5 filament assembly at internal dsRNA sites

Crystal structures of LGP2-RNA complexes identified dsRNA blunt ends as the preferred binding site for LGP2 (8–10) on dsRNA, but subsequent electron microscopy and atomic force microscopy studies suggested LGP2 can bind dsRNA internally as a stem binder (10,41). ATP hydrolysis by LGP2 has been reported to enhance binding of LGP2 to RNA and hence promote MDA5 signaling (39). To gain further insight on how LGP2 may affect MDA5 filament formation we imaged filaments formed in the presence of LGP2, MDA5 and dsRNA by ns-EM. The electron micrographs showed that addition of increasing amounts of LGP2 to mixtures of MDA5 and dsRNA progressively reduced the average length of the resulting protein filaments (**Figure 4A,B**). This could in principle be attributed to increased MDA5 filament nucleation, reduced cooperativity of MDA5 filament assembly, or competition between MDA5 and LGP2 for dsRNA binding. The overall ATPase activity of filaments and their capacity to displace streptavidin from biotinylated internal nucleotides were not significantly affected by LGP2 when the number of RNA binding sites was not limiting (**Figure 2D,E**), indicating that LGP2 did not interfere with the cooperativity of filament assembly or ATP hydrolysis by MDA5 filaments. Bilayer interferometry with immobilized dsRNA and a molar excess of 15-bp dsRNA binding sites showed that an MDA5-LGP2 mixture had a slightly higher net dsRNA binding affinity than MDA5 alone, with apparent *K_d_* values of 520 nM and 1.3 µM, respectively (**Figure 2B**). This suggests that the reduction in MDA5 filament length caused by LGP2 is primarily due to increased MDA5 filament nucleation rather than competition between MDA5 and LGP2 for dsRNA binding. However, when the number of dsRNA binding sites was limiting, the net ATPase activity of mixed MDA5-LGP2 filaments decreased with increasing concentrations of LGP2 (**Figure 4C**). Since the ATPase activity of LGP2 is one third of that of MDA5 (**Figure 2E**) and LGP2 has a higher binding affinity for dsRNA than MDA5 (**Figure 2B**), this decrease in ATPase activity can be attributed to competition between LGP2 and MDA5 for dsRNA binding sites. Whether MDA5 or LGP2 dsRNA binding sites are limiting in infected cells remains unknown but given that MDA5 binds RNA more cooperatively and is expressed at higher levels than LGP2, both at baseline and upon induction by viral infection or interferon (4,11), it seems unlikely that LGP2 would outcompete MDA5 for RNA binding in the cellular context. Overall, together with previous evidence that LGP2 amplifies MDA5 signaling (39–41), our data support the model that LGP2 promotes nucleation of MDA5 filaments – including at internal RNA sites with LGP2 ATP hydrolysis – resulting in shorter, more numerous filaments.

**Figure 4.**
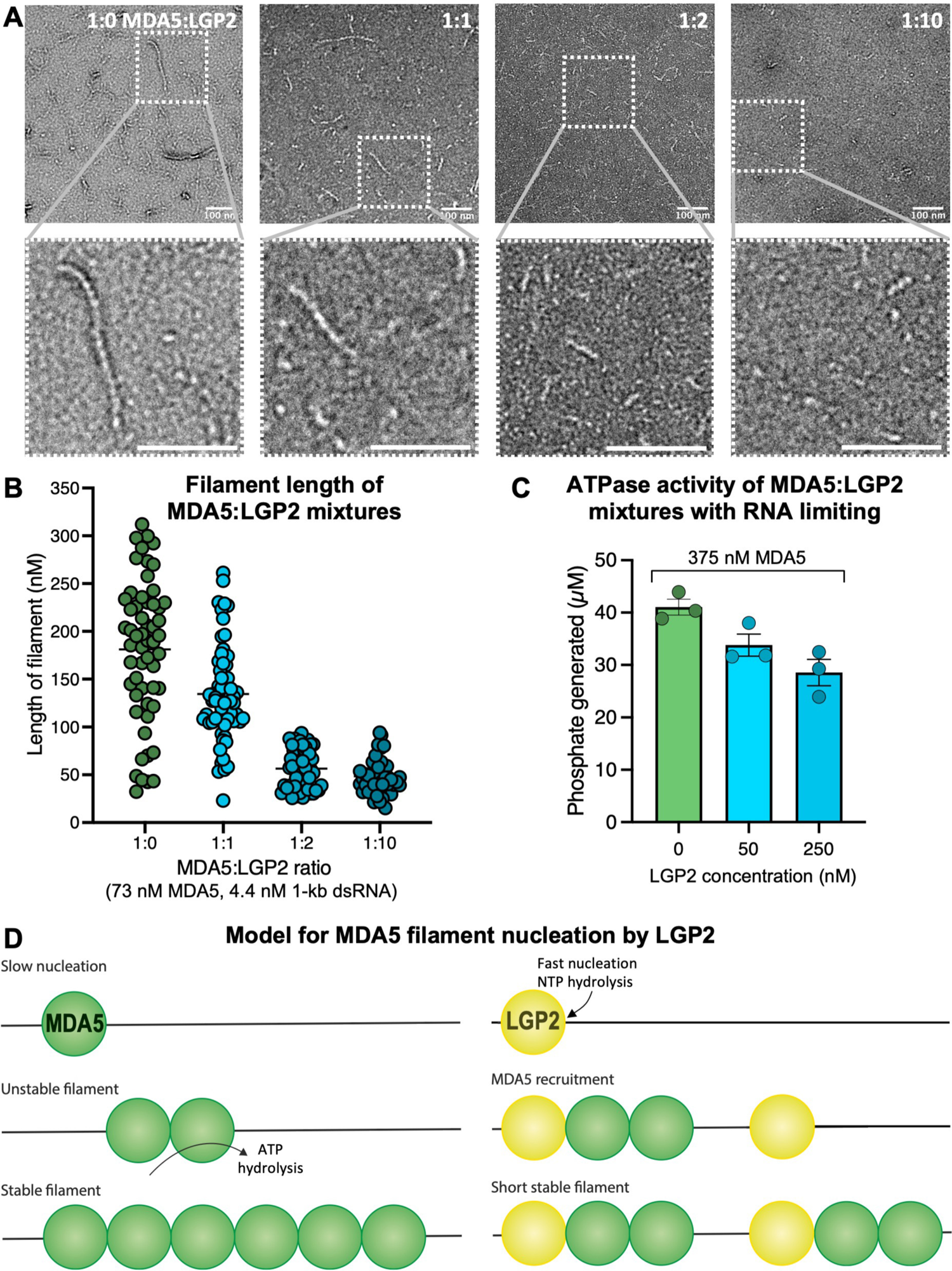
LGP2 promotes nucleation of MDA5 filament assembly at internal dsRNA sites. (A) Negative stain electron micrographs of filaments formed in the presence of 1-kb dsRNA and different proportions of MDA5 and LGP2. Scale bars, 100 nm. (B) Filament length measurements taken from electron micrographs including those shown in (A). Increasing amounts of LGP2 progressively reduced the average length of the resulting protein filaments. The number of measurements in each category from left to right was 56, 64, 53, and 48, respectively. See **Supplementary Data** for source data for panels B and C. (C) ATPase activity of filaments formed in the presence of MDA5, LGP2 and 1-kb dsRNA with a sub-stoichiometric number of 15-bp dsRNA binding sites. The concentration of 15-bp dsRNA binding sites was 313 nM. Error bars show the s.e.m. of three independent experiments. (D) Proposed NTPase-dependent mechanism of MDA5 filament nucleation by LGP2 at internal dsRNA sites.

### Molecular modeling of a LGP2-MDA5 heterodimer with AlphaFold-Multimer

Nucleation of MDA5 filament assembly by LGP2 may depend on direct interactions between MDA5 and LGP2. We could not confirm the presence of such interactions experimentally because the two proteins have similar structures and could not be distinguished in our electron micrographs. We therefore generated atomic model predictions of MDA5-LGP2 complexes with AlphaFold-Multimer (43–45) as implemented by ColabFold (45). With only the sequences of human or mouse LGP2 and MDA5 as the input, AlphaFold generated LGP2-MDA5 complexes with high confidence scores (average pLDDT = 85-87). All the generated AlphaFold models had the same overall structure, with or without the use of model templates from the Protein Data Bank. The predicted LGP2-MDA5 interface resembled the MDA5-MDA5 interface in cryo-EM structures of helical MDA5-dsRNA filaments (21). Despite RNA modeling not being supported by AlphaFold, the models contained a central void, or channel along the helical axis where dsRNA should bind (**Figure 5A,B**). In all models, LGP2 and MDA5 were in a head-to-tail configuration, with the MDA5 pincer domain at the center of the filament-like interface. When a second molecule of MDA5 was added to the modeling input, the same LGP2-MDA5 heterodimer was predicted and the second MDA5 molecule was placed on the opposing filament forming face of the first MDA5 molecule to form a three-subunit filament with LGP2 at one end (**Figure 5A**). Analysis with the PISA server (46) showed that the predicted LGP2-MDA5 interface had a larger buried surface area and estimated solvation free energy gain when compared to the MDA5-MDA5 interfaces obtained by cryo-EM or predicted by AlphaFold (**Figure 5C**).

**Figure 5.**
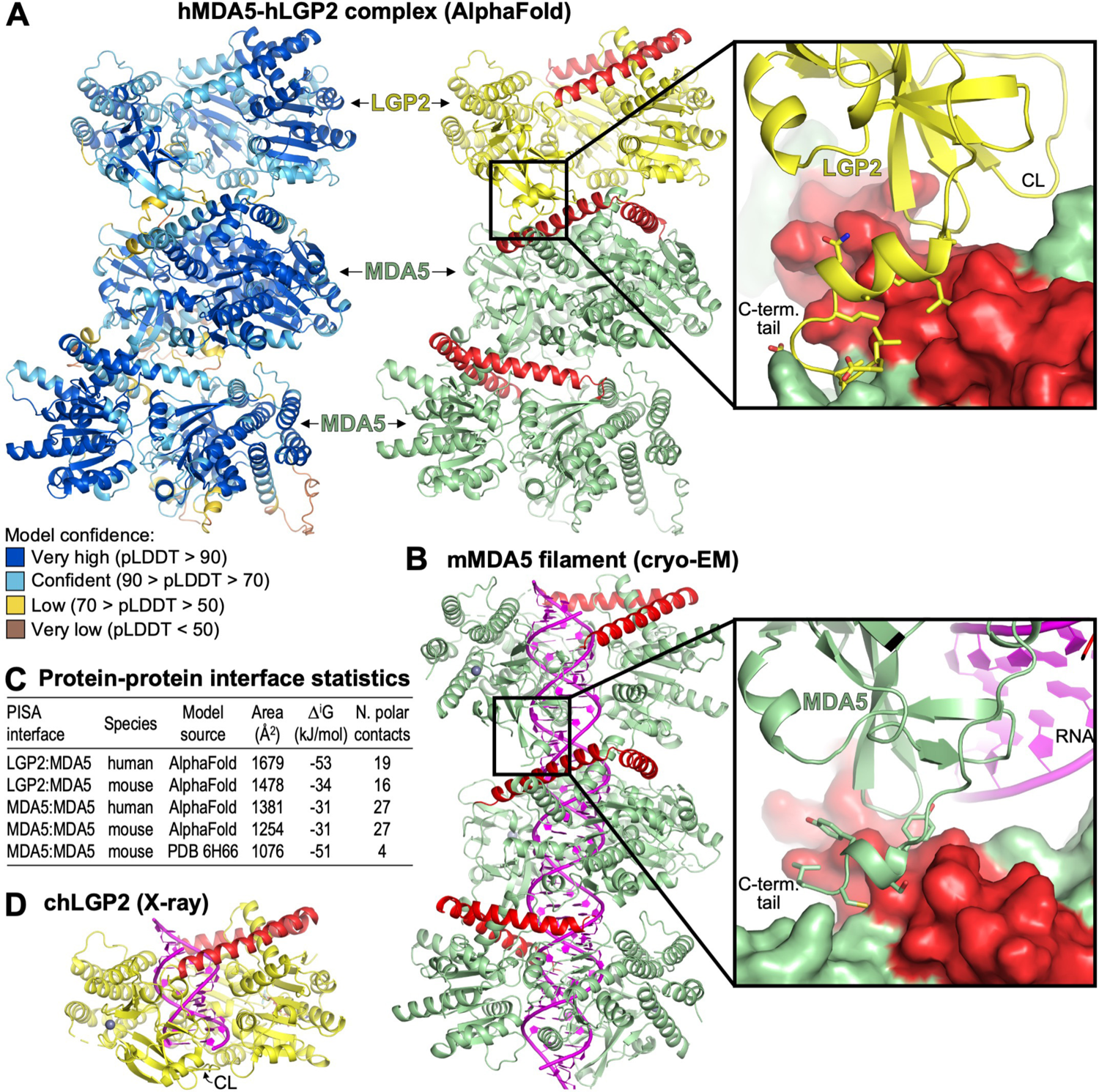
Molecular modeling of MDA5-LGP2 complexes with AlphaFold-Multimer. (A) AlphaFold model of a complex containing one LGP2 molecule and two MDA5 molecules colored by model confidence (pLDDT, according to the color code shown), left, and colored with MDA5 in green, LGP2 in yellow and the pincer domains of MDA5 and LG2 in red, right. The closeup shows the key contacts between LGP2 (yellow) and MDA5 (in surface representation). CL, capping loop. (B) Cryo-EM structure of the MDA5-dsRNA filament without nucleotide and with 93° helical twist (PDB 6H66 (21)). This cryo-EM structure was selected because it lacks nucleotide and has similar helical twist to the AlphaFold models. The dsRNA is shown in magenta. The closeup shows contacts between adjacent MDA5 protomers at the region equivalent to that shown in the closeup in (A). (C) Protein-protein interface statistics from PISA (46) for MDA5 filaments and MDA5-LGP2 complexes modeled by AlphaFold (43–45). Statistics for the cryo-EM-derived MDA5 filament shown in (B) are shown for comparison. (D) Crystal structure of chicken LGP2 in complex with ADP-AlF4-Mg and an end-bound 10-bp dsRNA (PDB 5JAJ (10)). CL, capping loop.

In crystal structures of LGP2-RNA complexes, LGP2 is in end-binding mode (10). The capping loop (residues 591-602) in the CTD caps one end of the RNA while allowing 5’ or 3’ overhangs to be accommodated (**Figure 5D**). Internal or stem binding to dsRNA would require the capping loop to move away from the helical axis of the dsRNA (10). In our AlphaFold models, the conformation of LGP2 is similar to that of the crystal structures, but LGP2 is unexpectedly oriented with its capping loop facing the interacting MDA5 subunit. This orientation means that the free end of an end-bound dsRNA would be located on the opposite side of LGP2, near the pincer domain and facing away from the MDA5 subunit. In the modeled conformation, the LGP2 capping loop partially occludes the dsRNA binding channel and would have to move away from the filament axis to accommodate dsRNA in the stem binding mode (**Figure 5A,B**). The polarity of the MDA5-LGP2 interaction – with the LGP2 C-terminal tail (residues 663-677) forming most of the contacts with MDA5 and the LGP2 pincer domain facing away from the interface – is the opposite of what would have been expected if LGP2 nucleated MDA5 filaments from dsRNA ends. Indeed, to bind the free dsRNA end in an end-bound LGP2-dsRNA complex, MDA5 would need to form contacts with the LGP2 pincer domain, on the side opposite to the capping loop and C-terminal tail.

## DISCUSSION

We have shown that adjacent MDA5 subunits in MDA5-dsRNA filaments hydrolyze ATP cooperatively. Since ATP hydrolysis promotes dissociation of MDA5 from suboptimal dsRNA ligands including endogenous RNA, cooperative ATP hydrolysis will trigger cooperative disassembly of MDA5 filaments form these ligands. Accordingly, the ATPase activity of mixtures of wild-type and ATPase-deficient MDA5 variants dropped exponentially as the amount of the ATPase-deficient variant was increased. Indeed, a 1:1 mixture of wild-type and ATPase-deficient MDA5 had only 20% of the ATPase activity of wild-type MDA5 (**Figure 1A**). Mutations targeting the ATPase activity of MDA5 can therefore be expected to have a dominant-negative effect on ATPase activity, and hence a dominant gain-of-function effect on MDA5-dependent interferon signaling activity. This provides a likely explanation for why ATPase-deficient MDA5 mutations are dominant autoinflammation alleles (2,32,33).

An additional consequence of cooperative ATP hydrolysis by MDA5 is to amplify the one base-pair RNA footprint expansion that accompanies each cycle of ATP hydrolysis by a single MDA5 molecule, allowing MDA5 filaments to displace bound viral or cellular proteins from the bound dsRNA. Nucleation of MDA5-dsRNA filament formation by LGP2 reduces MDA5 filament length and hence the cooperativity of MDA5 ATP hydrolysis, which should in turn increase filament stability. These mechanisms confer protein displacement and filament repair activities independent of interferon signaling. Despite these activities, however, we note that some virally encoded proteins cannot be displaced from RNA by MDA5, such as filovirus VP35 proteins, for example (52,53).

A-to-I RNA editing by ADAR1 prevents MDA5 signaling from endogenous RNAs by weakening base-pairing in double-stranded segments (27,28). In the absence of ADAR1 editing, endogenous dsRNAs trigger an MDA5-dependent interferon response that absolutely requires LGP2 (42). Cellular dsRNAs, such as inverted Alu repeats, are rarely longer than 150 bp and contain base-pair mismatches even without editing, making them poor ligands for MDA5. This suggests that LGP2, with its much lower nucleic acid selectivity, functions as an adaptor to nucleate MDA5 filament assembly on short or imperfect dsRNA ligands that would not otherwise support MDA5 signaling. LGP2 may have a similar adaptor, or intercellular surveillance function in initiating MDA5 signaling from virus-derived dsRNA ligands that would not otherwise activate MDA5 signaling, thereby broadening the repertoire of viral RNAs sensed by the cell (54).

Our ATPase assays show that ATP hydrolysis modulates the RNA binding affinities of MDA5 and LGP2 in contrasting manners. Whereas ATP hydrolysis by MDA5 promotes dissociation from suboptimal dsRNA ligands, ATP hydrolysis by LGP2 promotes binding to internal sites on dsRNA, in the stem binding mode. In this respect LGP2 is more similar to RIG-I, in which ATP hydrolysis also promotes binding to internal dsRNA binding sites (36,37). However, in contrast to RIG-I which reaches internal binding sites by local translocation from dsRNA ends (36,37), our data show that LGP2 can bind directly to internal sites on dsRNA.

Although helicases from other groups within superfamily 2 hydrolyze all four types of NTPs (49,50), the semi-selective NTP hydrolysis of LGP2 distinguishes it from MDA5. We speculate that LGP2 may have evolved to broaden its nucleotide usage as a mechanism to maintain NTPase-dependent dsRNA binding, MDA5 filament nucleation, and hence IFN-β activation when ATP is depleted or when GTP or CTP is elevated in the cell. Many RNA viruses have different genomic nucleotide compositions than their hosts. Retroviruses genomes are A-rich, for example, and the rubella virus genome has exceptionally high GC content (70%) (51). Infection with these viruses will increase utilization of specific nucleotides – GTP and CTP in the case of rubella virus – which may become depleted or subject to increased turnover. Indeed, HIV, rubella virus and Epstein-Barr virus all induce host cells to increase expression of rate-limiting enzymes in the nucleotide biosynthesis pathway (55–57).

Together with previous evidence that LGP2 promotes MDA5 signaling and filament nucleation (39–41), our ATPase and electron microscopy data support the model that LGP2 utilizes its NTPase activity to bind to dsRNA at internal binding sites, from where it nucleates MDA5 filament assembly. AlphaFold robustly predicts an MDA5-LGP2 interaction with LGP2 in an orientation that is incompatible with MDA5-LGP2 complex formation at dsRNA ends. Hence, the AlphaFold modeling suggests that LGP2 may not nucleate MDA5 filaments from dsRNA ends but rather from internal sites on dsRNA as a stem binder. However, this hypothesis remains to be tested experimentally.

We conclude that LGP2 promotes NTPase-dependent nucleation of MDA5 filaments at internal sites on dsRNA – and possibly NTPase-independent nucleation at dsRNA ends – resulting in shorter, more numerous filaments that are still capable of signaling. This work identifies novel molecular mechanisms contributing the selectivity and sensitivity of cytosolic dsRNA sensing by RIG-I-like receptors, with relevance to both healthy individuals and those with MDA5 mutations that result in autoinflammatory disease.

## DATA AVAILABILITY

The data underlying this article are available in the article and in its online supplementary material.

## SUPPLEMENTARY DATA

Supplementary Data are available at NAR online.

## AUTHOR CONTRIBUTIONS

Conceptualization, R.S. and Y.M.; Methodology, R.S., Y.W., A.H.d.V., K.E.L., T.H., B.F., and Y.M.; Investigation, R.S., Y.W., A.H.d.V., K.E.L., T.H., and M.C.; Formal Analysis, Y.M.; Writing – Original Draft, Y.M.; Writing – Review & Editing, R.S. and Y.M.; Visualization, R.S., Y.W., A.H.d.V, and Y.M.; Supervision, B.F. and Y.M.; Project Administration, B.F. and Y.M.; Funding Acquisition, R.S., A.H.d.V., B.F. and Y.M.

## ACKNOWLEDGEMENTS

We thank Giuseppe Cannone and Grigory Sharov (MRC-LMB EM Facility) for assistance in EM data collection. We thank Stephen McLaughlin and Chris Batters (MRC-LMB Biophysics Facility) for assistance with Octet experiments. We thank Irene Díaz-López (MRC-LMB) for providing the ssRNAs with complex secondary structure. We thank David Dulin (VU University Amsterdam) for providing the circular dsRNA. We thank Frank Adolf (University of Cambridge) for his advice on baculovirus expression. We thank members of the Modis lab for insightful discussions. We thank MRC-LMB Scientific Computing for computing support and Sami Chaaban (MRC-LMB) for cluster maintenance of ColabFold. We would like to acknowledge the support of the MRC-LMB Media & Glass Wash facility. We thank Jianguo Shi (MRC-LMB Baculovirus Facility) for assistance with insect cell culture.

