## Supplementary Figures for "Contrasting functions of ATP hydrolysis by MDA5 and LGP2 in viral RNA sensing"

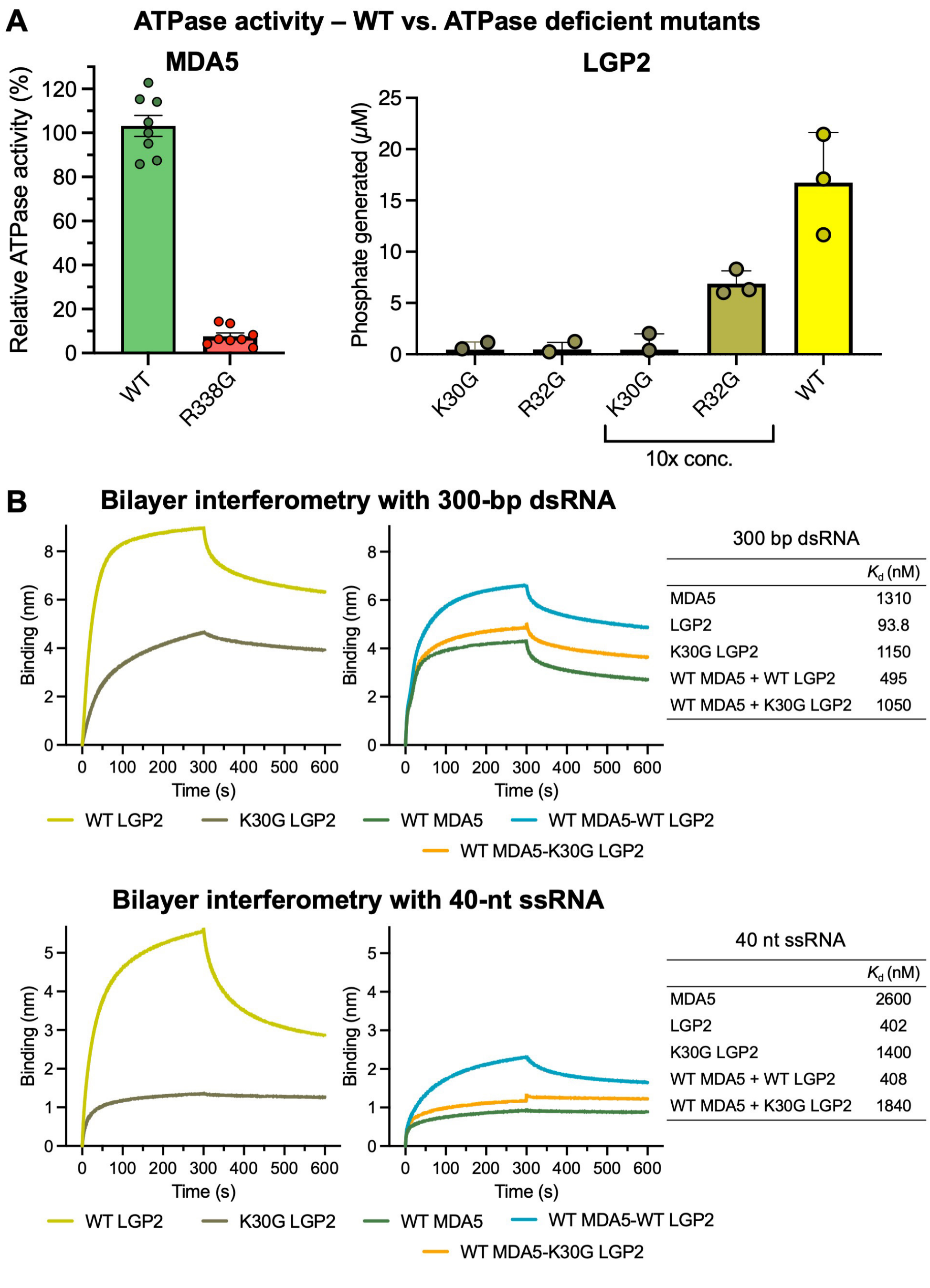


300 bp dsRNA

| | $K_d$ (nM) |
|---|---|
| MDA5 | 1310 |
| LGP2 | 93.8 |
| K30G LGP2 | 1150 |
| WT MDA5 + WT LGP2 | 495 |
| WT MDA5 + K30G LGP2 | 1050 |

40 nt ssRNA

| | $K_d$ (nM) |
|---|---|
| MDA5 | 2600 |
| LGP2 | 402 |
| K30G LGP2 | 1400 |
| WT MDA5 + WT LGP2 | 408 |
| WT MDA5 + K30G LGP2 | 1840 |

**Supplementary Figure S1.** ATPase and RNA binding assays with MDA5 and LGP2. (**A**) ATPase activities of R338G MDA5 (left), and K30G LGP2 and R32G LGP2 (right), in the presence of 1-kb dsRNA. Error bars show the s.e.m. of eight and three independent experiments for MDA5 and LGP2, respectively. (**B**) Bilayer interferometry with 3'-biotinylated 300-bp dsRNA (top) or 40-nt ssRNA (bottom) immobilized on a streptavidin biosensor and MDA5, LGP2 or MDA5-LGP2 mixtures in the mobile phase. Dissociation constants derived from the curves are tabulated.

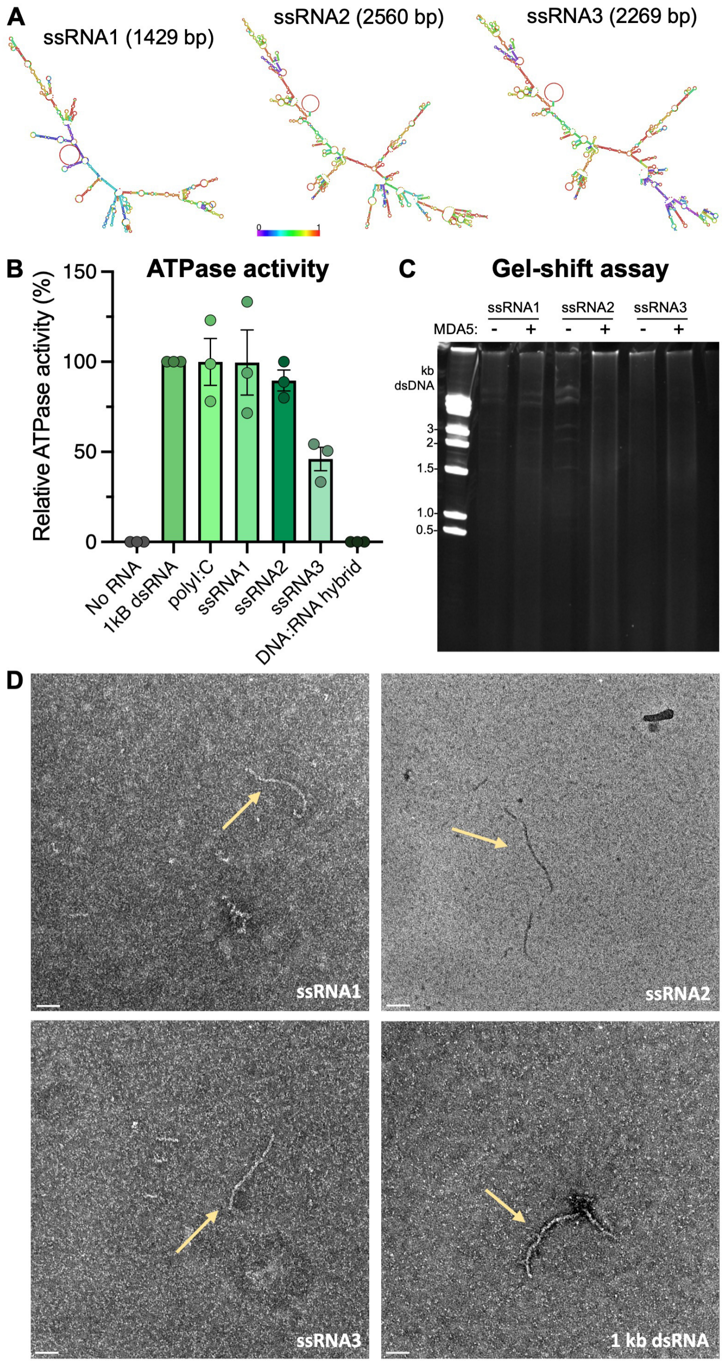


**Supplementary Figure S2.** ATPase and RNA binding assays for MDA5 and long ssRNAs with complex secondary structure. (**A**) Minimum Free energy (MFE) secondary structures of the ssRNAs used here, predicted by RNAfold [http://rna.tbi.univie.ac.at] and colored by base pairing probability using the color scale shown. (**B**) MDA5 ATPase activity in the presence of the RNAs shown in (A), along with high-molecular weight poly(I:C) and a 1-kb DNA:RNA hybrid, expressed as percent of ATPase activity with 1-kb dsRNA. Error bars show the s.e.m. of three independent experiments. (**C**) Polyacrylamide gel-shift assay for binding of MDA5 to the RNAs shown in (A). (**D**) Negative-stain electron micrographs of MDA5 filaments formed in the presence the RNAs shown in (A). Scale bars, 50 nm.
